## Supplemental Figs for "CRISPR/Cas9-based mutagenesis frequently provokes on-target mRNA misregulation"

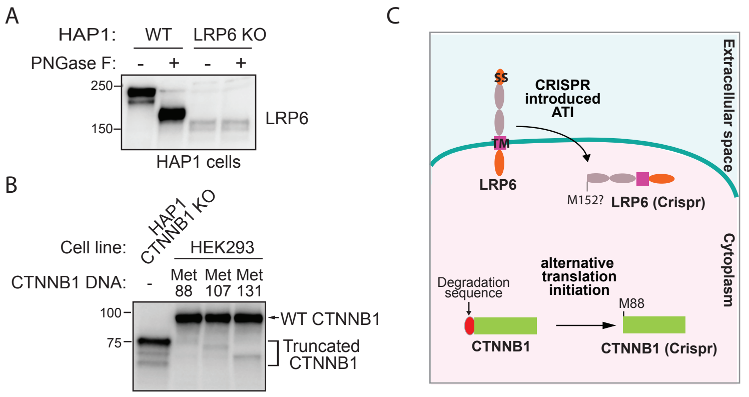


**Supplemental figure 1. Inadvertent production of novel proteins by CRISPR/Cas9. (A)** The LRP6 protein generated by ATI is no longer glycosylated. Lysates generated by HAP1 WT cells or with CRISPR/Cas9 editing in the LRP6 gene were incubated with deglycosidase PNGase F and subjected to Western blot analysis. **(B)** CRISPR/Cas9-edited cells express a CTNNB1 protein lacking the N-terminal GSK3β phosphorylation sites (Ser33 and Ser37) that promote β-TrCP and CTNNB1 protein turnover1. CTNNB1 cDNAs initiating at methionine 88, 107 or 131 were transiently transfected in HEK293 cells. CTNNB1 protein that initiates at methionine 88 co-migrated with the short CTNNB1 protein found in CRISPR-edited HAP1 cells. **(C)** CRISPR/Cas9 induced mislocalization of a type I transmembrane receptor. ATI results in bypassing of the signal sequence found in LRP6 WNT receptor, likely resulting in an intracellularly localized protein and produces CTNNB1 lacking degradation sequence.

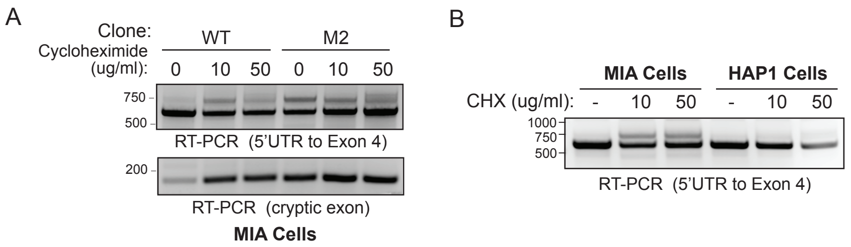


**Supplemental figure 2. CRISPR/Cas9-introduced INDELs facilitate the conversion of a pseudo-mRNA to a protein encoding mRNA. (A)** LKB1 pseudo-mRNA undergoes non-sense mediated decay (NMD) in MIA WT cells. MIA WT and clone M2 were incubated with 10 µg/ml or 50 µg /ml of NMD inhibitor cycloheximide (CHX). RT-PCR analysis using primers encompassing 5'UTR and exon 4 or the cryptic exon was performed. **(B)** Genomic sequence alteration by CRISPR/Cas9 gene editing must be superimposed on all the transcript variants native to different cell lines. RT-PCR using primers flanking 5' UTR and Exon 4 of LKB1 revealed an mRNA species that is sensitive to CHX in MIA cells, but not in HAP1 cells.

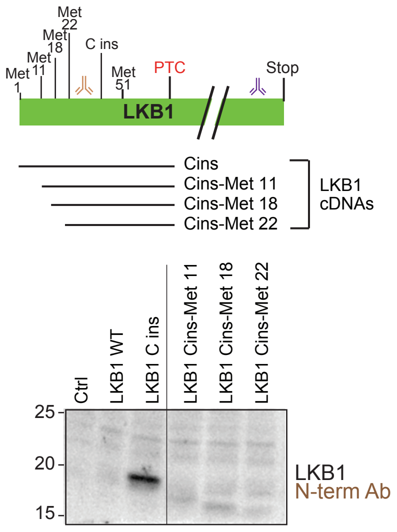


**Supplemental figure 3. Leaky scanning does not contribute to the CRISPR/Cas9-mediated production of novel proteins.** Overexpression constructs of *LKB1* gene with either WT sequence or C insertion initiating at methionine 1, 11, 18 or 22 were engineered and transiently transfected in Hela cells. Lysates generated from the cells were used to perform Western blot analysis with N-terminal LKB1 antibody.

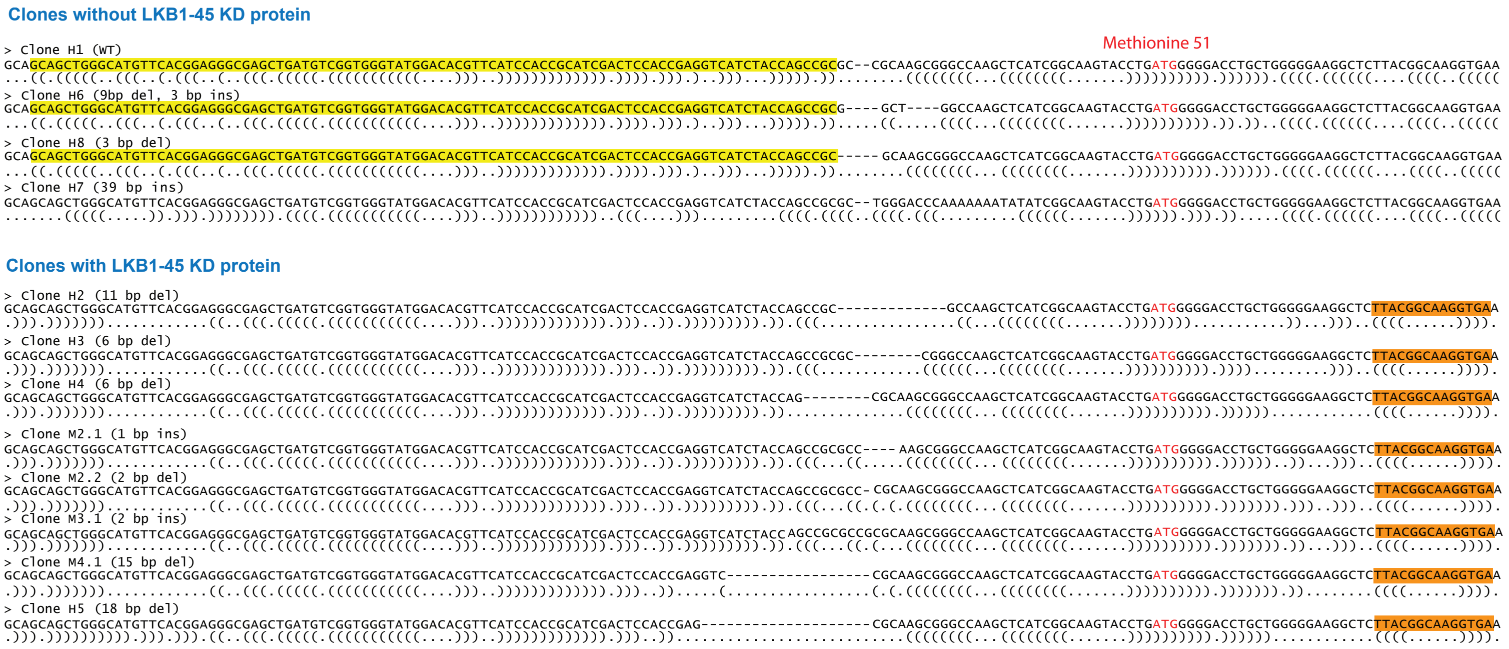


**Supplemental figure 4. RNA secondary structure analysis predicts motifs and features unique to CRISPR/Cas9-edited clones with ATI.** Turbofold II, an RNA structure prediction algorithm, was used to predict secondary structures for 12 different mutations in LKB1 generated by CRISPR/Cas9 gene editing using an sgRNA targeting exon 1 in MIA and HAP1 cells. Mutations in clones H1, H6, H7 and H8 were not associated with ATI. Mutations in clones H2, H3, H4, H5, M2.1, M2.2, M3.1, and M4.1 were associated with ATI. The predicted RNA structures are shown by dots and brackets along with alignments, where dot represents an unpaired nucleotide and bracket represents paired nucleotides. The structures of non-ATI clones H1, H6 and H8 show a unique helix (in yellow background) located before the Methionine 51 site (ATG highlighted in red). All eight sequences associated with ATI are predicted to form a conserved hairpin loop 3‘ to Methionine 51 start site (TTACGGCAAGGTGA, in orange).

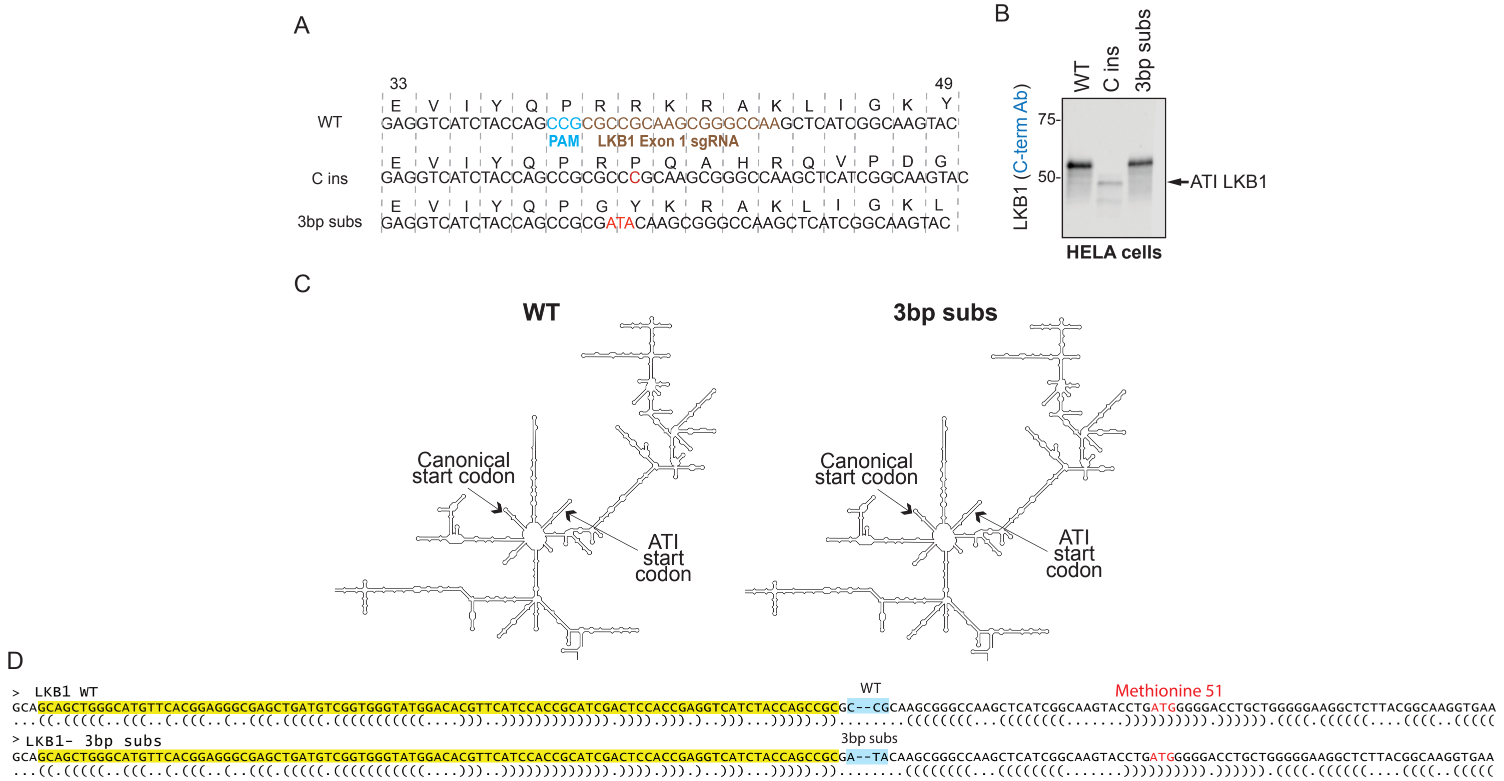


**Supplemental figure 5. Point mutagenesis does not replicate the ATI-inducing effect of INDELs introduced by CRISPR/Cas9. (A)** Expression plasmids with WT LKB1, the C insertion introduced by CRISPR/Cas9 in edited cell lines associated with ATI of LKB1, or a 3 bp substitution were engineered. **(B)** Lysates from HELA cells transiently transfected with LKB1 plasmids described in “A” were used to perform Western blot analysis with C-terminal LKB1 antibody. **(C)** No change in predicted RNA secondary structure was observed with the 3 bp substitution. **(D)** The predicted RNA structure represented by dot and brackets for WT LKB1 and LKB1 with the 3 bp substitution.

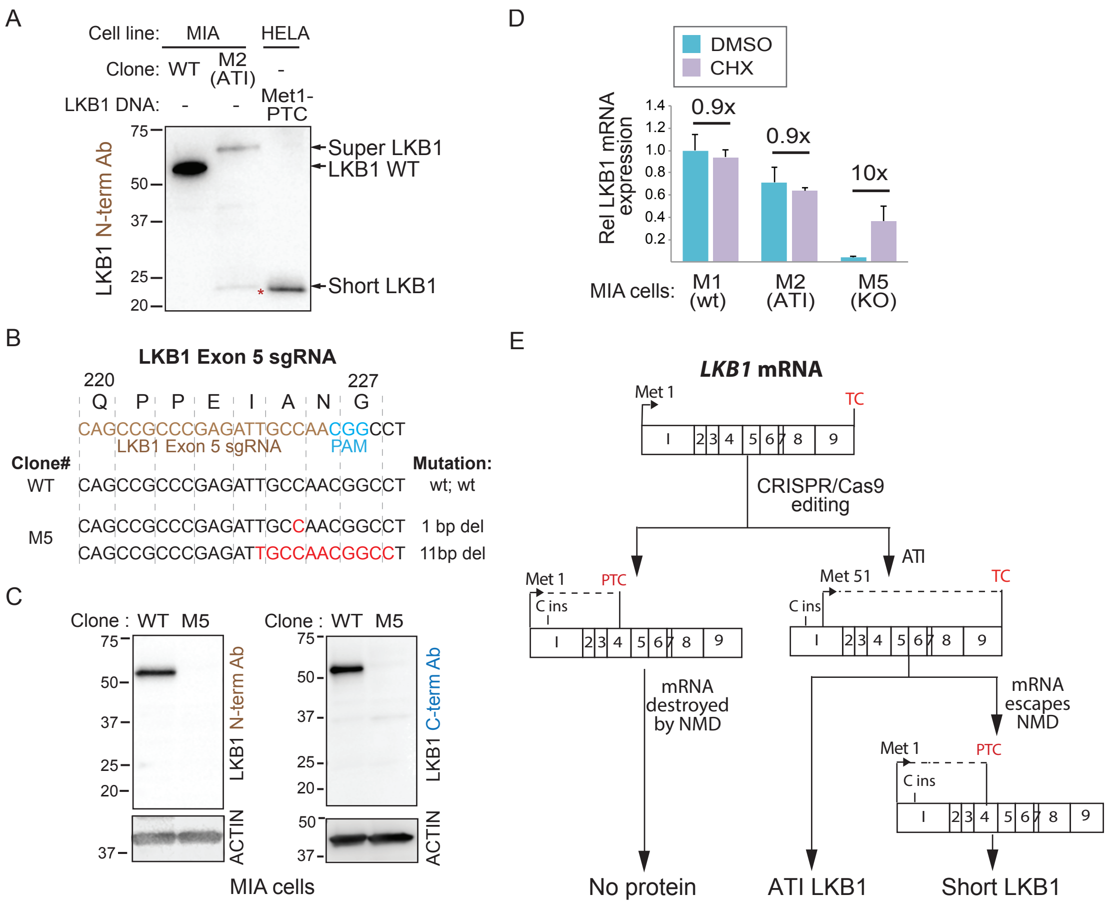


**Supplemental figure 6. ATI bypasses the induction of NMD in the presence of a pre-mature termination codon (PTC). (A)** CRISPR/-Cas9-edited MIA clones targeting LKB1 exhibiting ATI express a C-terminally truncated LKB1 protein (Short LKB1). Lysates from MIA LKB1 clones were subjected to Western blot analysis using an LKB1 antibody recognizing an N-terminus localized epitope. An LKB1 overexpression construct with canonical start site and the predicted PTC introduced by CRISPR/Cas9-editing was transiently transfected in HELA cells and analyzed along with the MIA clones. **(B)** Targeting an internal LKB1 exon as a strategy for generating MIA LKB1 KO cells. Genomic sequence of a MIA clone M5 edited with LKB1 exon 5 sgRNA indicates presence of a frameshift alteration. **(C)** Engineering MIA cells with no detectable expression of LKB1 proteins. Western blot analysis of MIA clone M5 was performed using LKB1 N- and C-terminal antibodies. **(D)** ATI prevents nonsense-medicated decay (NMD). MIA WT, clone M2 (ATI) and clone M5 (LKB1 KO) cells were treated with DMSO or 10 µg/ml CHX for 6 hrs and isolated cDNAs was subjected to quantitative RT-PCR (qPCR) using primers flanking Exons 3 and 4. The MIA clone with a non-ATI inducing mutation in LKB1 exhibits a 10-fold change in mRNA abundance upon CHX treatment. CHX treatment has no effect on LKB1 mRNA abundance in cells with ATI. **(E)** Production of novel LKB1 proteins as a consequence of a CRISPR/Cas9-induced INDEL.

| Genes | Targeted exon | CRISPR-introduced INDELs | INDELs hit ESEs? | Exon skipped? |
|---|---|---|---|---|
| CTNNB1 | 3 | 4bps del | No | No |
| AXIN1 | 2 | 1 bps ins | No | No |
| LRP6 | 2 | 5 bps del | No | No |
| TBK1 | 1 | 2 bps ins | No | No |
| BAP1 | 5 | 109 bps ins | No | No |
| TLE3 | 7 | 2 bps del | No | Yes |
| PPM1A | 1 | 5 bps del | No | No |
| BCL2L2 | 1 | 5 bps del | No | No |
| SUFU | 3 | 11bps del, 1bp ins | No | No |
| SUFU | 8 | 1 bps del | No | No |
| RICTOR | 5 | 10 bps del | Yes | No |
| VPS35 | 5 | 11 bps del | Yes | No |
| TOP1 | 6 | 8 bps del | Yes | Yes |
| SIRT1 | 4 | 20 bps del | Yes | Yes |
| PTEN | 1 | 5 bps del | Yes | No |
| SUFU | 2 | 5 bps del, 11 bp del | Yes | Yes |
| SUFU | 3 | 26 bps del, 2 bps del | Yes | Yes |
| SUFU | 8 | 28 bps del, 1 bp ins | Yes | Yes |
| SUFU | 8 | 11 bp del | Yes | Yes |
| SUFU | 8 | 2 bps del | Yes | Yes |
| SUFU | 8 | 1 bp del | Yes | Yes |
| SUFU | 8 | 46 bps ins | Yes | Yes |
| SUFU | 8 | 61 bps ins | Yes | Yes |
| SUFU | 8 | 1 bp ins | Yes | Yes |

**% Exon skipped when ESE is compromised: 78% (11/14)**

**% Exon skipped when ESE is not compromised: 10% (1/10)**

**Supplemental figure 7. CRISPR/Cas9-edited cells with different mutations analyzed for ESE disruption and exon skipping.** List of 24 CRISPR/Cas9-edited cells from Horizon Discovery and *de novo* engineering with indicated targeted exons, frameshift inducing mutations, predicted impact on ESEs, and exon skipping status derived from RT-PCR analysis.

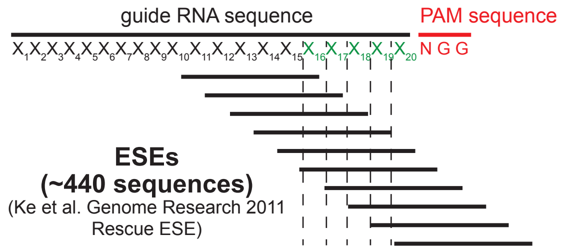


**Supplemental figure 8. ESE sequence evaluation using the CRISPinatoR.** The CRISPinatoR algorithm selects for sgRNAs containing ESEs within 5bp 5' to the PAM sequence. 440 putative ESE sequences previously described (http://genes.mit.edu/burgelab/rescue-ese/ and Ke et al.) were used to develop the CRISPinatoR algorithm.

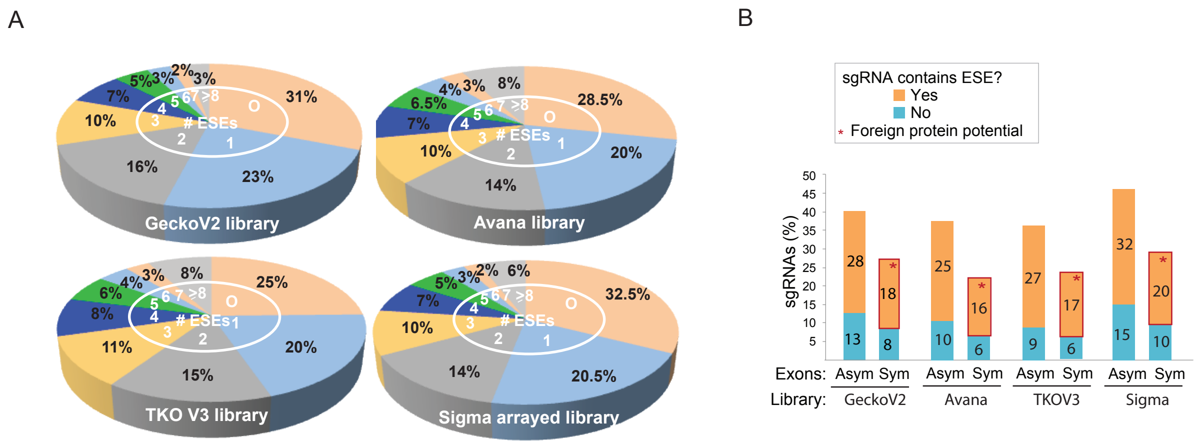


**Supplemental figure 9. ESE sequence targeting prevalence in genome-wide CRISPR/Cas9 libraries. (A)** ESE annotation in commercial CRISPR libraries. Putative ESEs embedded within the sgRNA sequences present in 4 different CRISPR libraries were analyzed: Gecko V2, Avana, TKO V3, and Sanger. **(B)** Genome-wide CRISPR/Cas9 screening libraries do not avoid inadvertent targeting of ESEs. Exon symmetry and ESE targeting potential were calculated for the different CRISPR/Cas9 libraries. sgRNAs with putative ESEs targeting symmetric exons (red box) have the potential to generate foreign proteins.

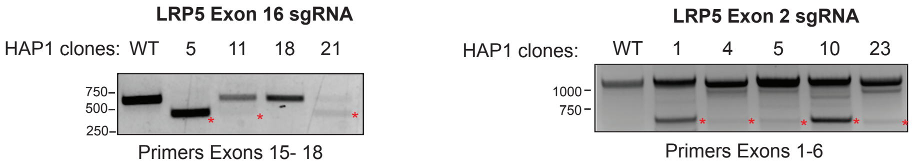


**Supplemental figure 10. Induction of exon skipping with targeted disruption of ESEs in an asymmetric or symmetric exon of LRP5. (A)** RT-PCR analysis of HAP1 clones edited with the LRP5 Exon 16 sgRNA. **(B)** RT-PCR analysis of HAP1 clones edited with the LRP5 Exon 2 sgRNA.

**Supplemental References**


1. Winston, J.T. et al. The SCFbeta-TRCP-ubiquitin ligase complex associates specifically with phosphorylated destruction motifs in IkappaBalpha and beta-catenin and stimulates IkappaBalpha ubiquitination in vitro. *Genes Dev* **13**, 270-283. (1999).
2. Ke, S. & Chasin, L.A. Context-dependent splicing regulation: exon definition, co-occurring motif pairs and tissue specificity. *RNA Biol* **8**, 384-388 (2011).
3. Sanjana, N.E., Shalem, O. & Zhang, F. Improved vectors and genome-wide libraries for CRISPR screening. *Nature methods* **11**, 783-784 (2014).
4. Doench, J.G. et al. Optimized sgRNA design to maximize activity and minimize off-target effects of CRISPR-Cas9. *Nature biotechnology* **34**, 184-191 (2016).
5. Hart, T. et al. Evaluation and Design of Genome-Wide CRISPR/SpCas9 Knockout Screens. *G3 (Bethesda)* **7**, 2719-2727 (2017).
6. Metzakopian, E. et al. Enhancing the genome editing toolbox: genome wide CRISPR arrayed libraries. *Sci Rep* **7**, 2244 (2017).
